# Induced neural progenitor cells and iPS-neurons from major depressive disorder patients show altered bioenergetics and electrophysiological properties

**DOI:** 10.1101/2021.04.30.441774

**Authors:** Julian Triebelhorn, Iseline Cardon, Kerstin Kuffner, Stefanie Bader, Tatjana Jahner, Katrin Meindl, Tanja Rothhammer-Hampl, Markus J. Riemenschneider, Konstantin Drexler, Mark Berneburg, Caroline Nothdurfter, André Manook, Christoph Brochhausen, Thomas C. Baghai, Sven Hilbert, Rainer Rupprecht, Vladimir M. Milenkovic, Christian H. Wetzel

## Abstract

The molecular pathomechanisms of major depressive disorder (MDD) are still not completely understood. Here, we follow the hypothesis, that mitochondria dysfunction which is inevitably associated with bioenergetic disbalance is a risk factor that contributes to the susceptibility of an individual to develop MDD. Thus, we investigated molecular mechanisms related to mitochondrial function in induced neuronal progenitor cells (NPCs) which were reprogrammed from fibroblasts of eight MDD patients and eight non-depressed controls. We found significantly lower maximal respiration rates, altered cytosolic basal calcium levels, and smaller soma size in NPCs derived from MDD patients. These findings are partially consistent with our earlier observations in MDD patient-derived fibroblasts. Furthermore, we differentiated MDD and control NPCs into iPS-neurons and analysed their passive biophysical and active electrophysiological properties to investigate whether neuronal function can be related to altered mitochondrial activity and bioenergetics. Interestingly, MDD patient-derived iPS-neurons showed significantly lower membrane capacitance, a less hyperpolarized membrane potential, increased Na^+^ current density and increased spontaneous electrical activity. Our findings indicate that functional differences evident in fibroblasts derived from MDD patients are partially present after reprogramming to induced-NPCs, might relate to altered function of iPS-neurons and thus might be associated with the aetiology of major depressive disorder.

## INTRODUCTION

Depressive syndrome is a debilitating and severe mental disorder, affecting about 5 million people per year in Germany and 350 million people worldwide (World Health Organization, 2016). The molecular mechanisms underlying the cause and progress of this complex disease are still not completely understood. Hypotheses argue for a combination of neurobiological factors which affect cellular function and neuronal communication, thereby increasing the risk for MDD, in conjunction with environmental (psychosocial) stress. The interaction of all factors has been shown to be associated with the onset and severity of depressive episodes^1^. Involved molecular mechanisms include genetically inherited and acquired neurobiological issues such as the dysregulation of monoaminergic, glutamatergic and GABAergic neurotransmission as well as reduced neuroplasticity and neurogenesis as a consequence of impaired BDNF signaling (reviewed in^2^). Furthermore, mitochondrial dysfunction associated with reduced bioenergetic capability is considered an important risk factor for MDD as well as other psychiatric disorders^3-5^, by constraining proper cellular function and rendering the cells vulnerable to stress, especially during increased metabolic demand. Mitochondrial disorders that present with MDD as well as mitochondrial abnormalities and involved pathomechanisms in patients living with depression are reported in ^6-10^ and reviewed in ^3, 5, 11, 12^.

Especially brain neurons rely heavily on mitochondrial energy provision to maintain membrane excitability and operate neurotransmission, as well as to sustain plasticity^13^. However, MDD is increasingly viewed as a systemic disease with somatic manifestations outside the brain^10, 14^. Literature already reports mitochondria-related effects in fibroblasts^6, 10^, muscle-^9^, peripheral mononuclear blood cells^7^, and platelets^8, 15^ of depressed patients, suggesting that mitochondria-related pathomechanisms associated with MDD can be identified and studied in neuronal and also in peripheral non-neuronal cells.

To challenge the hypothesis that mitochondria dysfunction contributes to the aetiology of the disease, we aimed to identify and characterize molecular pathomechanisms related to mitochondrial function and bioenergetic imbalance in a human cellular model of major depression^16-19^. We used skin fibroblasts from eight MDD patients and eight non-depressed controls^6^ to reprogram and differentiate these cell lines to neural progenitor cells (NPCs) and neurons (iPS-neurons), shifting our cellular model from non-neuronal to neuronal cells. The advent of induced pluripotent stem cell (iPSC) technology^20, 21^ opens a new avenue for studying basic biological and pathological mechanisms^17, 22, 23^. The generation of disease- and patient-specific iPSCs and their differentiation into specific cell types helps unravelling the molecular mechanisms, which underlie the etiology of complex diseases such as MDD^16, 17^. As a measure of mitochondrial function in MDD and non-depressed control NPCs, we assessed the performance of the oxidative phosphorylating system (OXPHOS) by analysing the oxygen consumption rates in different respiratory states and the mitochondrial membrane potential (MMP) as an indicator for the bioenergetic state. Moreover, the cellular ATP content was analysed as the primary energetic outcome of mitochondrial function. The cytosolic Ca^2+^ level provided important information about cellular Ca^2+^ homeostasis, since Ca^2+^ ions serve as important signalling molecules regulating most of mitochondrial and cellular functions, ranging from electron transport chain (ETC) to apoptosis. Interestingly, the reprogrammed NPCs presented functional alterations, which might be associated with bioenergetic dysfunction of mitochondria in MDD patient cells and were in line with our findings obtained in the founder fibroblasts^6^.

Moreover, to characterize electrophysiological properties of neurons derived from MDD patients and to identify molecular phenotypes of mitochondrial impairment in these cells, we differentiated the NPCs into iPS-neurons. iPS-neurons were analysed by means of whole-cell voltage- and current-clamp recordings. We found differences in the membrane capacitance, resting membrane potential, Na^+^ current density and spontaneous activity of MDD iPS-neurons when compared with cells derived from non-depressed controls. In summary, we identified alterations in bioenergetic parameters in NPCs, and found altered electrophysiological properties in iPS-neurons of patients suffering from depression, which might be related to reduced bioenergetics. Moreover, our findings indicate that functional differences evident in fibroblasts from MDD patients^6^ are partially present after reprogramming and differentiation to induced-NPCs, thereby affecting functional properties of iPS-neurons. We discuss that the underlying mechanisms might be associated with the aetiology of major depressive disorder. Mitochondrial alterations or altered energy metabolism in iPSC-derived neural lineages has already been studied^24-30^, but to the best of our knowledge, this is the first report addressing altered energy metabolism in iPSC-derived neural lineages in the context of major depression.

## METHODS AND MATERIALS

### Skin Biopsies and primary human fibroblast cultivation

Skin biopsies were conducted by the Department of Dermatology, University Hospital of Regensburg, Regensburg, Germany. All participants gave written informed consent, and all study procedures were approved by the ethics committee of the University of Regensburg (ref: 13-101-0271). Human fibroblasts were obtained and cultivated as previously described^6^. Until use, cells were stored in a robotic storage system (SmartFreezer®, Angelantoni) in the Central Biobank Regensburg at the vapor phase of liquid nitrogen and standardized conditions.

### Generation of control and MDD patient iPSCs

iPSCs were generated using the episomal protocol described by^21^. Briefly, 5×10^5^ fibroblasts of passage 4 or less were electroporated with 600 ng of each the episomal vectors pCBX-EBNS, pCE-hsk, pCE-hUL, pCE-hOCT3/4 and pCE-mp53DD using the Amaxa Nucleofactor (Lonza). The cells were then cultured in TeSR-E7 medium for 3-4 weeks on Matrigel-coated dishes (Corning) until the colonies appeared. iPSC colonies were manually picked and cultured using mTeSR1 medium on Matrigel (StemCell Technologies). The generated iPSC cultures were analysed for pluripotency by the PluriTest® method, a patented bioinformatics assay for the quality assessment of iPSCs, which was adjusted for next-generation sequencing (NGS) data as published by^31^.

### iPSC differentiation to NPCs and neuron differentiation

NPCs were derived from iPSCs using a protocol described in^32^. Briefly, NPCs were expanded in expansion medium containing 50% Neurobasal medium, 50% Advanced DMEM/F12, and neural induction supplement, (all from Life Technologies), and subsequently dissociated with Accutase (Life Technologies) and maintained on Geltrex-coated dishes at a density of 1×10^5^ cells per cm^2^. To obtain mature neurons, NPCs from passages 5 to 10 were plated onto poly-L-ornithine/laminin (Sigma-Aldrich) coated glass cover-slips and differentiated in Neurobasal medium supplemented for 21 days with 1% B27, 0.5% GlutaMax, 0.5% non-essential amino acids, 0.5% Culture One (Thermo Fisher Scientific), 200 nM ascorbic acid (Carl Roth), 20 ng/ml BDNF, 20 ng/ml GDNF (both PeproTech), 1 mM dibutyryl-cAMP (Stemcell), 4 μg/ml laminin (Sigma) and 50 U/ml Penicillin, 50 μg/ml Streptomycin (Thermo Fisher Scientific).

### Immunofluorescence

Human fibroblasts, iPSCs, NPCs and induced neurons grown on glass coverslips (Menzel Gläser), were fixed in 4% paraformaldehyde (Carl Roth) and blocked by 10% normal goat serum in PBST (PBS+0.5% Triton X-100) for 20 min. Cells were then incubated with primary antibodies in PBST (PBS+0.1% Triton X-100, 2% normal goat serum) overnight at 4 °C, following 1h incubation at RT with corresponding Alexa 488/Cy3/Cy5 coupled secondary antibodies (1:1000, Thermo Fisher Scientific). Cell nuclei were stained with DAPI. Fibroblasts were labelled with anti-SMA (mouse; 1:250, ab7817, Abcam), whereas iPSCs were immunostained using anti-NANOG (rabbit; 1:250, ab21624, Abcam) antibodies. NPCs were labeled with anti-PAX6 (mouse; 1:10; deposited to the DSHB by Kawakami, A.) and anti-SOX2, ab97959 antibodies (rabbit; 1:1000, ab97959, Abcam). For immunostaining of the induced neurons following antibodies were used: anti-Tuj 1 (mouse; 1:2000, G7121, Promega), anti-MAP2 (chicken; 1:5000, ab5392, Abcam), anti-synaptophysin (rabbit; 1:500, ab52636, Abcam), anti-VGLUT1 (rabbit; 1:1000, ab180188, Abcam), and anti-NEUN (rabbit; 1:500, ab177487, Abcam). Coverslips were fixed onto glass object slides (Menzel Gläser) using Fluorescence Mounting Medium (Dako) and images were acquired using a Zeiss Observer Z.1 microscope (Zeiss). For quantitative analysis of immunofluorescence labelling, all nuclei (DAPI) of NPCs of three visual fields (inspected with a 40X objective) were counted and the % of cells, that were/were not stained/double stained with PAX6 and SOX2 were calculated.

### qPCR analysis of expression of neuronal markers

Total RNA was isolated from iPSCs and from induced neurons after 21 DIV using RNA Plus Kit (Macherey-Nagel) according to the manufacturer’s instructions. First strand cDNA synthesis from 1 µg of total RNA was performed with QuantiTect Reverse Transcription Kit (Qiagen). Quantitative RT-PCR experiments were performed with Rotor-Gene-Q machine (Qiagen) using the 1x Takyon SYBR Master Mix (Eurogentec) and intron-spanning primers specific for different neural subtypes, listed in Supplementary Table 1. Measurements were completed in triplicate and results were analyzed with a Rotor-Gene-Q software version 2.3 (Qiagen). Relative expression levels of each target gene were calculated using the comparative C_t_ method and HPRT1 as a housekeeping gene for normalization. Gene expression of various neural subtypes was represented as heat map without statistical analysis.

**Table 1.** PluriTest results from all cells used in this study.

| Sample | Result | Pluripotency (>1424) | Novelty (<2.5) |
| --- | --- | --- | --- |
| CON1 | Pluripotent | 1540.3 | 1.99 |
| MDD1 | Not Pluripotent | 1336.6 | 2.32 |
| CON2 | Not Pluripotent | 1403.1 | 2.44 |
| MDD2 | Pluripotent | 1513.4 | 2.45 |
| CON3 | Pluripotent | 1426.4 | 2.49 |
| MDD3 | Pluripotent | 1473.5 | 2.34 |
| CON4 | Not Pluripotent | 1392.2 | 2.41 |
| MDD4 | Pluripotent | 1598.5 | 2.25 |
| CON5 | Pluripotent | 1442.1 | 2.24 |
| MDD5 | Pluripotent | 1524.2 | 2.20 |
| CON6 | Pluripotent | 1457.6 | 2.27 |
| MDD6 | Pluripotent | 1488.8 | 2.31 |
| CON7 | Pluripotent | 1597.9 | 2.33 |
| MDD7 | Pluripotent | 1504.7 | 2.38 |
| CON8 | Pluripotent | 1521.2 | 2.49 |
| MDD8 | Pluripotent | 1661.4 | 2.32 |

### Analysis of mitochondrial respiration

Analysis of mitochondrial respiration was performed using Seahorse XFp Flux analyzer with a Seahorse XFp Mito Stress Test Kit (Agilent Technologies) according to manufacturer’s recommendations. Briefly, 8×10^4^ NPCs were grown in XFp 8-well miniplates at 37 °C, humidified air and 5% CO_2_. Oxygen consumption rate (OCR) was measured with sequential injection of 1 µM Oligomycin, 1 µM FCCP, and each 0.5 µM Rotenone/Antimycin A (Biomol). After the measurements, nuclei were stained with DAPI and counted in order to to normalize respiration data.

### Luminescent Assay for ATP content

1×10^6^ cells were pelleted in a 1.5 ml Eppendorf cup and stored at -20 °C. ATP content was measured by CellTiter-Glo®Cell Viability Kit (Promega). CellTiter-Glo®Reagent containing CellTiter®Substrate and CellTiter®Buffer was thawed on ice. The ATP standard curve ranged between concentrations of 10 µM, 1µM, 100 nM, 10 nM and 1 nM ATP in 1x PBS. Cell pellets were resuspended in 500 µl PBS, heated at 100 °C for 2 min and kept on ice afterwards. Triplicates of 50 µl per sample and each standard were applied to a black 96-well-plate together with 50 µl of CellTiter-Glo®Reagent. The absorption was measured with the VarioScan at an integration time of 1 sec. The RLU generated by the SkanIT Software was used to calculate the actual ATP concentrations using the ATP standard curve. ATP concentrations were normalized to µg/mL protein using a BCA assay (Thermo Fisher Scientific).

### Mitochondrial membrane potential (JC-1)

Mitochondrial membrane potential was measured in NPCs as described in^33^. Briefly, 2.8×10^6^ neural progenitor cells were grown overnight on glass coverslips (diameter 25 mm; Menzel Gläser) and were loaded with 1 µM of JC-1/Pluronic in OptiMEM (Thermo Fisher Scientific) for 30 min at 37 °C (humidified air and 5% CO_2_). Mitochondrial membrane potential was compared as ratio of red over green fluorescence intensity in regions of interest, manually selected in the visual field using FIJI/ImageJ^34^. The macro used for the analysis will be provided on request.

### Imaging of cytosolic Ca^2+^ (Fura-2/AM)

Cytosolic Ca^2+^ levels in NPCs were measured as previously described in^35^. Briefly, 2.8×10^6^ neural progenitor cells were grown on glass coverslips in 6-well plate and loaded with 2 µM Fura-2/AM and Pluronic F127 in OptiMEM (Thermo Fisher Scientific) for 30 min at 37 °C. Cytosolic Ca^2+^ level in NPCs was assessed by measuring the fluorescence at 510 nm after excitation at 340 or 380 nm using Observer Z.1 inverted microscope (Zeiss). Cells were selected as regions of interest in the visual field using the FIJI/ImageJ^36^. The macro used for the analysis will be provided on request.

### Electrophysiology

Whole-cell patch-clamp recordings were performed on induced neurons during their 4th week of differentiation. A total of 333 induced neurons (162 Control-Neurons and 171 MDD-Neurons) were recorded and analysed. The extracellular solution was composed of 140 mM NaCl, 5 mM KCl, 2 mM CaCl_2_, 1 mM MgCl_2_, 10 mM HEPES, pH 7.3. Micropipettes were made of borosilicate glass (Science Products) by means of a horizontal pipette puller (Zeitz Instruments) and fire-polished to obtain a series resistance of 3-5 MΩ. Micropipettes were filled with intracellular solution (140 mM KCl, 1 mM MgCl_2_, 0.1 mM CaCl_2_, 5 mM EGTA, 10 mM HEPES). Recordings were made using a Heka Electronic EPC-10 amplifier (HEKA Electronic). The liquid-liquid junction potential was in the range of 12 to 16 mV and was corrected. The series resistance was assessed but not compensated. The resting membrane potential and capacitance were recorded directly after reaching the whole-cell configuration. For voltage-clamp recordings, membrane potential was held at -80 mV and depolarized in steps of 10 mV to evoke voltage-activated Na^+^- and K^+^-channels. In current-clamp mode, manually adjusted currents in the range of –2 to –240 pA (dependent on the resting membrane potential and the series resistance), were injected to hyperpolarize the membrane potential of neurons to about -80 mV. The cells were then depolarized by 20 steps with a 2 to 20 pA increment to reach the threshold and evoke action potentials. Spontaneous action potentials were recorded at a holding potential of about - 45 to -50 mV. All patch-clamp recordings were carried out at room temperature. Data were analyzed using Patchmaster v2×90.03. Cells with resting membrane potential of 0 mV and above were excluded from the analysis.

### Statistical Analysis

Graphical depiction and statistical analysis were conducted with Graph Pad Prism 8.0.2 (GraphPad Software). To exclude any bias, cell lines were renamed and anonymized before starting the differentiation protocol and experiments. For all experiments, except patch-clamp recordings, the means of two to three technical replicates were calculated and two to three biological replicates were averaged. Measurements were conducted pairwise allowing direct comparison of MDD vs. controls. Data were checked for normal distribution and appropriate statistical tests were applied (paired *t*-test, Wilcoxon matched-pairs signed rank test). Statistical outliers were detected and eliminated using ROUT-Method. Normal distribution was implied because of high sample size. Results of spontaneous activity were compared by Fisher’s exact test. Membrane potential showed no variance homogeneity, thus the Welch-test was applied. All other results showed variance homogeneity and were compared using unpaired Student’s *t*-Test. Results are presented as mean ± SEM, unless otherwise stated. *p*-value limit for statistical significance is set to ≤ 0.05.

Multilevel regression analyses were conducted using the statistical software R (Team RC, 2021). Alterations in the variables between pre- and posttests were analyzed using linear mixed regression models using the package “lme4”^37^. Type-I-Error probabilities for the regression coefficients were calculated using the package “lmerTest”^38^. Tests were run two-tailed. A detailed description of the model is given by ^39^.

## RESULTS

Based on our findings of altered bioenergetic properties and mitochondrial function in primary dermal fibroblasts of patients suffering from major depressive disorder (MDD)^6^, we were interested in the identification and characterization of MDD-associated pathomechanisms in neural cells. Thus, we used primary skin fibroblasts characterized in our earlier study^6^ (information on study participants is given in the Supplementary Table 2) and reprogrammed them to induced pluripotent stem cells (iPSCs) by transient episomal transduction according to the Yamanaka protocol (Figure 1A)^20, 21^. Pluripotency of iPSCs was validated by PluriTest analysis, a bioinformatics assay for the quality assessment of iPSCs by transcription profiling based on next-generation sequencing data^31^ obtained from the respective iPSC clones. The PluriTest analysis revealed that 13 of the iPSC clones used in this study could be considered pluripotent. Three others were very close to the empirical threshold (pluripotency >1424; novelty <2.5) (Figure 1B and Table 1). The iPSC colonies were differentiated to early neural progenitor cells (NPCs) within 7 days after culture medium was changed from mTeSR to neural induction medium. Further maturation of NPCs was allowed for 5 passages before cells were stained for neural progenitor markers SOX2 and PAX6^40^ (Figure 1C). Table 2 presents the percentage of cells positively labelled with the respective antibody and indicates that the majority of the cells co-expresses both NPC markers and can be regarded as neural progenitors. The three cell lines that failed the PluriTest analysis, showed prominent expression of the neuronal progenitor markers PAX6 and SOX2 as well, thus we decided to not exclude these cell lines from our study. To seek for functional differences between NPCs from MDD and non-depressed control subjects, we analysed bioenergetic properties and mitochondrial function.

**Table 2.** Generation of PAX6/SOX2 positive NPCs from human iPSCs via

| Cell line | PAX6 positive (%) | SOX2 positive (%) |
| --- | --- | --- |
| CON1 | 87.6 | 86.0 |
| MDD1 | 90.8 | 96.3 |
| CON2 | 90.7 | 90.0 |
| MDD2 | 82.2 | 85.4 |
| CON3 | 84.6 | 85.6 |
| MDD3 | 91.7 | 91.4 |
| CON4 | 86.4 | 90.8 |
| MDD4 | 84.3 | 86.9 |
| CON5 | 90.4 | 93.0 |
| MDD5 | 87.8 | 89.6 |
| CON6 | 93.1 | 92.9 |
| MDD6 | 91.8 | 89.2 |
| CON7 | 91.7 | 91.7 |
| MDD7 | 89.1 | 90.4 |
| CON8 | 88.6 | 89.7 |
| MDD8 | 88.3 | 90.2 |

**Figure 1.**
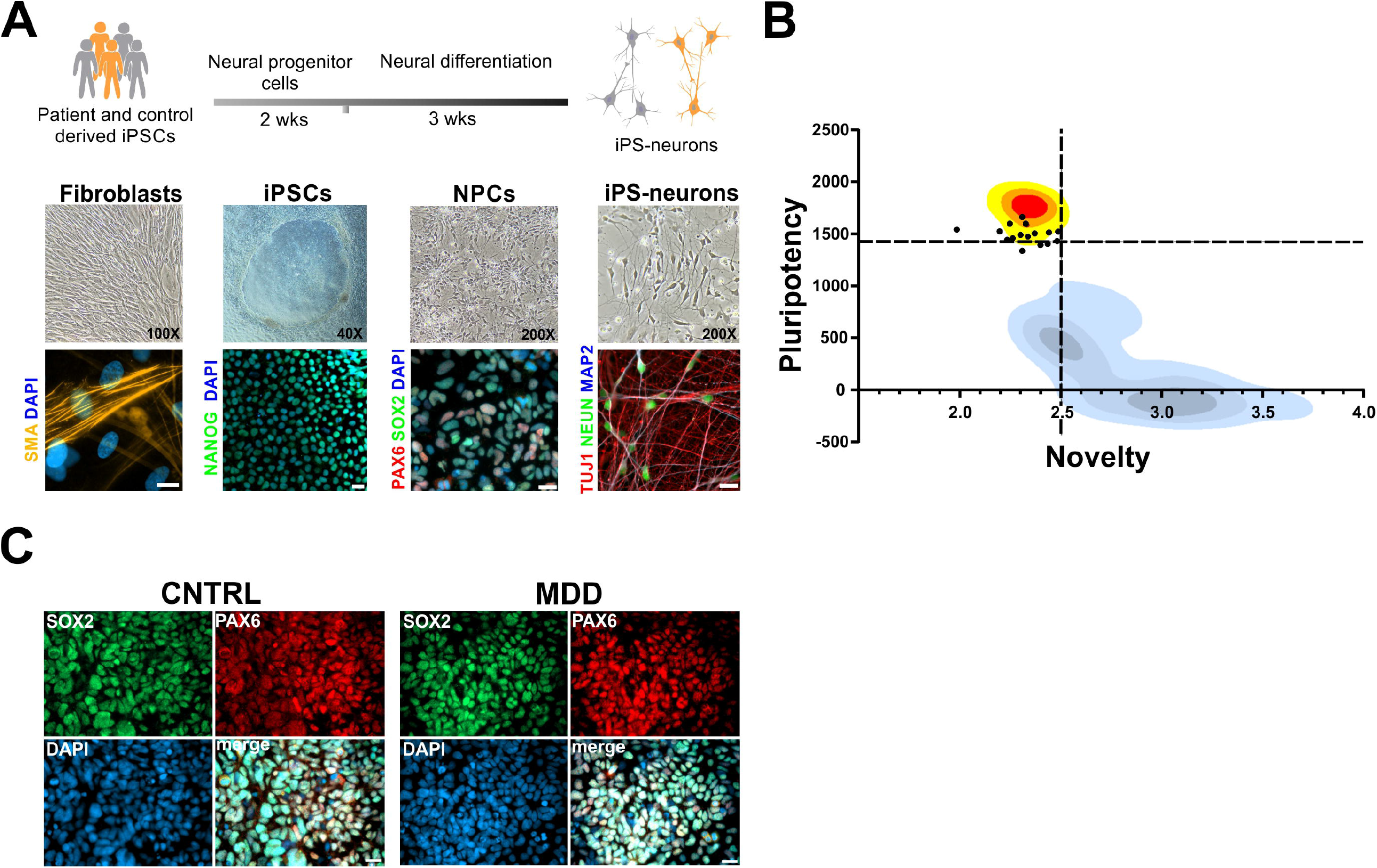
Generation of iPSC-based model of MDD. **(A)** Patient-derived dermal fibroblasts were reprogrammed into iPSCs and subsequently differentiated into neurons. Representative images of fibroblasts, iPSCs, NPCs and iPS-neurons expressing corresponding cell specific markers. Scale bar indicates 20 µm. **(B)** Characterization of 16 pluripotent cell lines (black dots) from this study using PluriTest. Kernel density estimations for iPSCs (yellow to red), and non-pluripotent stem cells (shades of blue/grey) are shown. Black dashed lines indicate the empirical thresholds for novelty and pluripotency. High quality cells should be located in the upper left quadrant, with a pluripotency score above the threshold and a novelty score below the threshold. It can be seen that the empirical thresholds detect high quality pluripotent cells from this study, but that this population still contains some cells with reduced pluripotency. **(C)** Representative images of NPCs differentiated from control and MDD patients show that majority of the cells are expressing typical neural progenitor cell markers PAX6 and SOX2. Scale bar indicates 20 µm.

### Mitochondrial membrane potential

The cationic and lipophilic fluorescent dye JC-1 accumulates in the mitochondrial membrane to an extent, which is dependent on the strength of the electric field. In negatively charged (i.e., highly energized) mitochondria, JC-1 molecules accumulate and form red fluorescing aggregates, while the fluorescence changes to green, when the dye molecules disaggregate into monomers in response to dissipation of the transmembrane potential. The ratio of the fluorescence signals emitted by the two states of JC-1 is then analyzed as a measure of the MMP. We found that NPCs from MDD patients presented a lower, however a non-significantly different red/green ratio when compared to the control NPCs from non-depressed subjects. This can be judged as an indication to a more depolarized MMP in patient cells (MDD 2.171 ± 0.16 vs. Cntrl 2.417 ± 0.114, mean ± SEM, *p* = 0.148, *t* = 21.291, *df* = 14.058, linear mixed regression) (Figure 2A).

**Figure 2.**
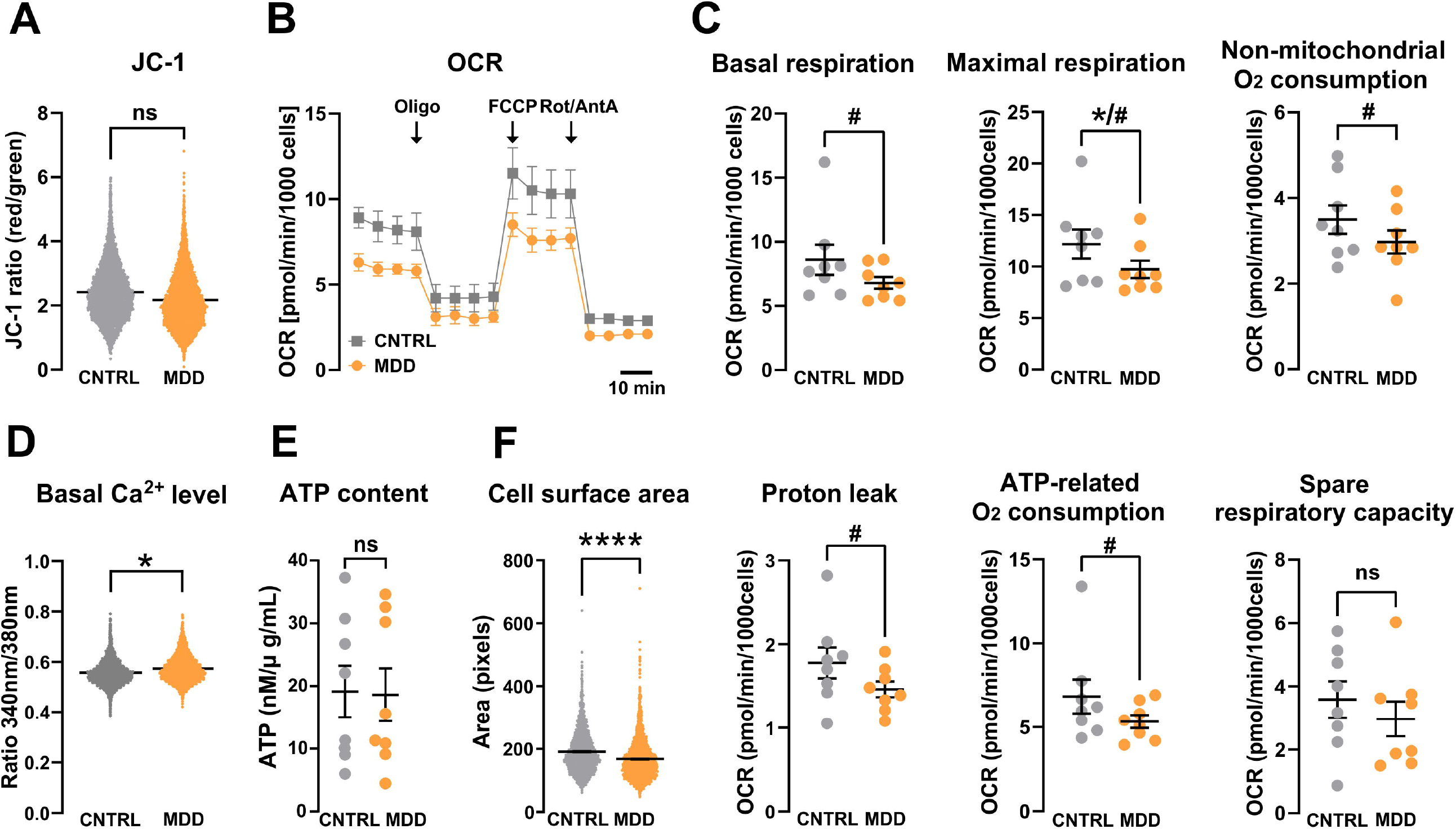
Mitochondrial bioenergetics. **(A)** Mitochondrial membrane potential of NPCs. Red/green (JC-1 aggregate/monomer) ratios of control (CNTRL) and MDD NPCs. Dot plots show mean red/green ratios ± SEM, control n=8, MDD n=8. Difference is not significant. **(B and C)** Oxygen consumption rates (OCR) of control and MDD NPCs. **(B)** Representative OCR measurement during the Mito Stress Test for MDD and control NPCs. **(C)** Maximal respiration is significantly different as analyzed by two-tailed *t*-test (indicated by *). Considering our hypothesis of reduced oxygen consumption in MDD cells, a one-tailed *t*-test indicate significant difference of all parameters except spare-respiratory capacity (indicated by #). Dot plots show normalized mean OCR values ± SEM, Control n=8, MDD n=8. (**D**) Basal Ca^2+^ calcium levels were significantly altered in MDD NPCs. Shown are the Fura-2 340 nm/380 nm fluorescence ratios of non-depressed control and MDD NPC lines. Dots show mean ratios (340 nm/380 nm, ratios ± SEM). Control n=8, MDD n=8. (**E**) ATP levels were not significantly different between MDD and CNTRL NPCs. Dots show normalized mean RLU values ± SEM. (**F**) MDD NPCs are significantly smaller than CNTRLs. Size is analyzed by assessing area (pixels) of Fura-2-loaded cells. Significant differences between MDD and non-depressive controls are indicated with *.

### Mitochondrial oxidative phosphorylation system (OXPHOS) - Respirometry

The consumption of molecular oxygen by accepting the electrons delivered by the ETC at complex IV can be used as a readout for the function and performance of the OXPHOS. Figure 2B depicts a representative Seahorse measurement using the Mito Stress Kit reporting the oxygen consumption rates (OCR) in various respiratory states. Although all assessed parameters were lower in patients suffering from depression, we found a significant difference between the groups in the maximal respiration only ([pmol/min/1000 cells] MDD 9.74 ± 0.85 vs. Cntrl 12.19 ± 1.41, *p* = 0.038, *t* = 2,551, *df* = 7, paired *t*-test, two-tailed) (Figure 2C). The basal respiration, as well as ATP-related oxygen consumption, non-mitochondrial OCR and proton leak were not significantly lower in MDD NPCs compared to healthy controls. However, based on our earlier results on the reduced respiration of fibroblasts from patients suffering from depression, we would expect/assume a reduced respiration in NPCs as well. Following this hypothesis, we can justify to use a one-tailed test instead, leading to significant differences between the MDD and control groups in the other respiratory parameters (Table 3).

**Table 3.** Statistical parameters of mitochondrial respiration

|  | Basal respiration |  | Maximal respiration |  | Non-mitochondrial OC |  | Proton leak |  | ATP-related OC |  | Spare respiratory capacity |  |
| --- | --- | --- | --- | --- | --- | --- | --- | --- | --- | --- | --- | --- |
| Group | CNTRL | MDD | CNTRL | MDD | CNTRL | MDD | CNTRL | MDD | CNTRL | MDD | CNTRL | MDD |
| n | 8 | 8 | 8 | 8 | 8 | 8 | 8 | 8 | 8 | 8 | 8 | 8 |
| Mean | 8.606 | 6.794 | 12.19 | 9.736 | 3.496 | 2.973 | 1.778 | 1.461 | 6.829 | 5.333 | 3.583 | 2.971 |
| Std. Error of Mean | 1.176 | 0.4575 | 1.413 | 0.8457 | 0.3333 | 0.271 | 0.1834 | 0.09506 | 1.011 | 0.3817 | 0.5777 | 0.5504 |
| *p* value two-tailed | 0.0879 |  | 0.038 |  | 0.0555 |  | 0.0585 |  | 0.0981 |  | 0.4081 |  |
| *p* value summary two-tailed | ns |  | * |  | ns |  | ns |  | ns |  | ns |  |
| *p* value one-tailed | 0.044 |  | 0.019 |  | 0.0277 |  | 0.0293 |  | 0.049 |  | 0.2041 |  |
| *p* value summary one-tailed | # |  | # |  | # |  | # |  | # |  | ns |  |
| *t* | 1.982 |  | 2.551 |  | 2.294 |  | 2.258 |  | 1.908 |  | 0.8798 |  |
| *df* | 7 |  | 7 |  | 7 |  | 7 |  | 7 |  | 7 |  |

### Cytosolic Ca^2+^ levels, cellular ATP-content, and cell size

As an additional measure for metabolic/bioenergetic activity and mitochondrial function, we compared the cytosolic Ca^2+^ levels in MDD and control NPCs by Fura-2 Ca^2+^ imaging via linear multilevel regression models, accounting for the nesting of multiple measurements within cell lines. We detected a small but significant difference in cytosolic Ca^2+^ levels between MDD and control NPCs (Fura-2 emission ratio: MDD 0.574 ± 0.008, Cntrl 0.558 ± 0.005, mean ± SEM, *t* = 2.247, *df* = 13.889, *p* = 0.041), consistent with the important function of Ca^2+^ as a signaling molecule and mediator in mitochondrial and cellular physiology (Figure 2D). Although we found reduced respiration and altered Ca^2+^ levels in MDD NPCs, the cellular ATP content was not different between the two groups ([nM/µg/ml] Cntrl 19.15 ± 4.11, MDD 18.62 ± 4.22, mean ± SEM, *p* = 0.69, *t*-test, two-tailed) (Figure 2E). This observation points to a compensatory mechanism in the NPCs, which compensates for a possibly reduced energy production according to a reduced OXPHOS. Although we could not detect a deficit in cellular ATP levels in the MDD NPCs, this group presented a significantly smaller cell size, as assessed by pixel counting of the fluorescent area during Fura-2 imaging ([pixel] MDD 168.1 ± 1.21, Cntrl 191.3 ± 1.29, mean ± SEM, *p* = 0.0001, *t*-test, two-tailed) (Figure 2F).

### Functional characteristics of iPS-neurons

Next, we were interested whether differential functional phenotypes can also be associated with iPS-neurons from MDD or control subjects. To this end, we differentiated NPC lines from eight MDD patients and matching controls to iPS-neurons. iPS-neurons showed clear bipolar or multipolar neuronal morphology (Figure 3A) and expressed the typical neuronal markers class III beta-tubulin, MAP2 and NEUN (Figure 3B). Moreover, they were immunopositive for the specific synaptic proteins VGLUT1 and SYP at DIV 21 (Figure 3C and D). A qPCR analysis of neuronal marker expression in 5 pairs of iPS-neuron lines indicated the expression of MAP2, GAD1, and GRIN1, whereas SLC6A4, TH, and CHAT showed only low expression (Figure 3E), indicating the differentiation of NPCs to functional neurons capable of glutamatergic and GABAergic signaling. The Ct values for NPCs and iPS-neurons are presented in the Supplementary Table 3. To functionally characterize and compare electrophysiological properties of iPS-neurons, we performed whole-cell voltage-clamp and current-clamp recordings of iPS-neurons during their 4^th^ week of differentiation and analyzed their passive biophysical as well as their active neuronal properties. Figure 4A-D shows exemplary whole-cell recordings of a typical iPS-neuron. In total, we recorded 333 neurons derived and differentiated from the eight MDD patients and eight non-depressed controls. We found that the membrane capacitance of iPS-neurons from MDD patients, which was compensated as C-slow by the inbuilt capacitance compensation algorithm of the EPC 10 amplifier, was significantly smaller compared with non-depressed control neurons ([pF] MDD 15.75 ± 0.76 vs. Cntrl 18.12 ± 0.9, *p* = 0.044, *t* = 2.022, *df* = 328, *t*-test, unpaired) (Figure 5A). Since C-slow is associated with the size of the membrane surface, the capacitance can be regarded as a measure for cell size. Interestingly, our data indicate that iPS-neurons from MDD patients are smaller than from non-depressed controls. This finding is consistent with the reduced cell size in MDD NPCs (Figure 2F). The series resistance was not different between the two groups ([MΩ] MDD 23.09 ± 1.035 vs. Cntrl 24.79 ± 1.11, *p* = 0.26, *t* = 1.121, *df* = 318, *t*-test, unpaired) (Figure 5B). Interestingly, we found that the resting membrane potential measured directly after establishing the whole-cell configuration was different between the groups: MDD iPS-neurons had a significantly lower (i.e., less hyperpolarized) resting membrane potential than iPS-neurons derived from the control subjects ([mV] MDD –23.68 ± 0.84 vs. Cntrl - 26.68 ± 1.02, *p* = 0.024, *t* = 2.275, *df* = 312.3, *t*-test, unpaired, with Welch’s correction) (Figure 5C). In current-clamp mode, the neurons were able to fire action potentials (APs) when the membrane potential was adjusted to approximately -80 mV by current injection and then depolarized by modulating the injected current. The mean current injected to adjust the membrane potential to -80 mV was not different between MDD and non-depressed control iPS-neurons ([pA] MDD -84.08 ± 5.214 vs. Cntrl –81.14 ± 5.29, *p* = 0.69, *t* = 0.40, *df* = 299, *t*-test, unpaired) (Figure 5D). In voltage-clamp experiments, voltage-activated sodium inward currents and potassium outward currents were induced by stepwise depolarization of the membrane potential starting at a V_hold_ of -80 mV. To consider the between-group difference in membrane capacitance as an indicator for different cell size, we analyzed the current density (pA/pF) by relating the Na^+^ and K^+^ current amplitudes to the measured membrane capacitance and found a higher I_Na+_ current density in MDD iPS-neurons ([pA/pF] MDD -47.4 ± 2.9 vs. Cntrl –35.2 ± 1.9, *p* = 0.0006, *t* = 3.465, *df* = 291, *t*-test, unpaired) (Figure 5E). The amplitudes of the potassium (I_K+_) outward currents at +20 mV were not different between MDD and non-depressed iPS-neurons ([pA/pF] MDD -40.8 ± 2.2 vs. Cntrl –41.9 ± 2.3, *p* = 0.7151, *t* = 0.3653, *df* = 307, *t*-test, unpaired) (Figure 5G). Assessing the amplitudes of I_Na+_ at 0 mV to minimize effects of intrinsic space clamp problems in large neurons, we again detected significantly larger I_Na+_ current density in MDD iPS-neurons ([pA/pF] MDD -35.15 ± 2.3 vs. Cntrl –27.09 ± 1.4, *p* = 0.0001, *t* = 11.71, *df* = 135, *t*-test, unpaired) (Figure 5F). In addition, we characterized spontaneous electrical activity of cultured iPS-neurons by current-clamp recording and adjusted the resting membrane potential to about -50 mV by direct current injection. Under this condition, we found that a significantly higher proportion of MDD-derived iPS-neurons (35/163 = 21.74%), were spontaneously active and generated APs when compared with cells reprogrammed from non-depressed controls (18/162 = 11.11%) indicating an increased activity of MDD neurons (*p* = 0.016, Fisher’s exact test) (Figure 5H).

**Figure 3.**
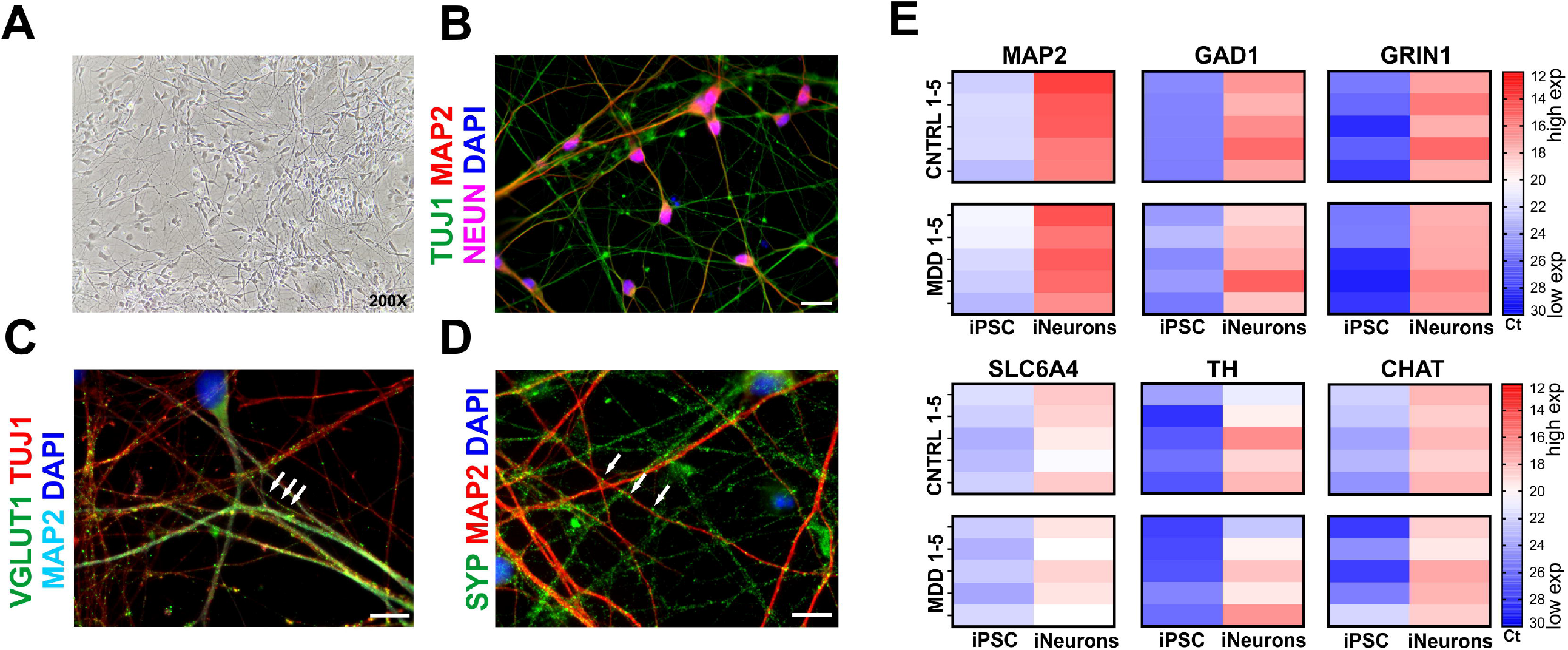
Characterization of iPS-neurons. **(A)** Neurons differentiated from NPCs show clear bipolar or -multipolar neuronal morphology. **(B)** Representative images of iPS-neurons expressing typical neuronal markers such as class III beta-tubulin (TUJ1), microtubule associated protein 2 (MAP2), and neuronal nuclear protein (NEUN). Scale bar indicates 20 µm. **(C, D)** Formation of glutamatergic synapses in iPS-neurons after 3-4 weeks in the culture. VGLUT1 and synaptophysin (SYP) immunostaining visualizes synaptic vesicle clusters (green, white arrows). Scale bar indicates 10 µm. **(E)** Heat map representation of relative gene expression by cycle threshold of RT-PCR reactions. Changes of expression of induced neurons after 21 days of neural differentiation relative to corresponding iPSCs are represented as downregulation (blue), no change (white) or upregulation (red).

**Figure 4.**
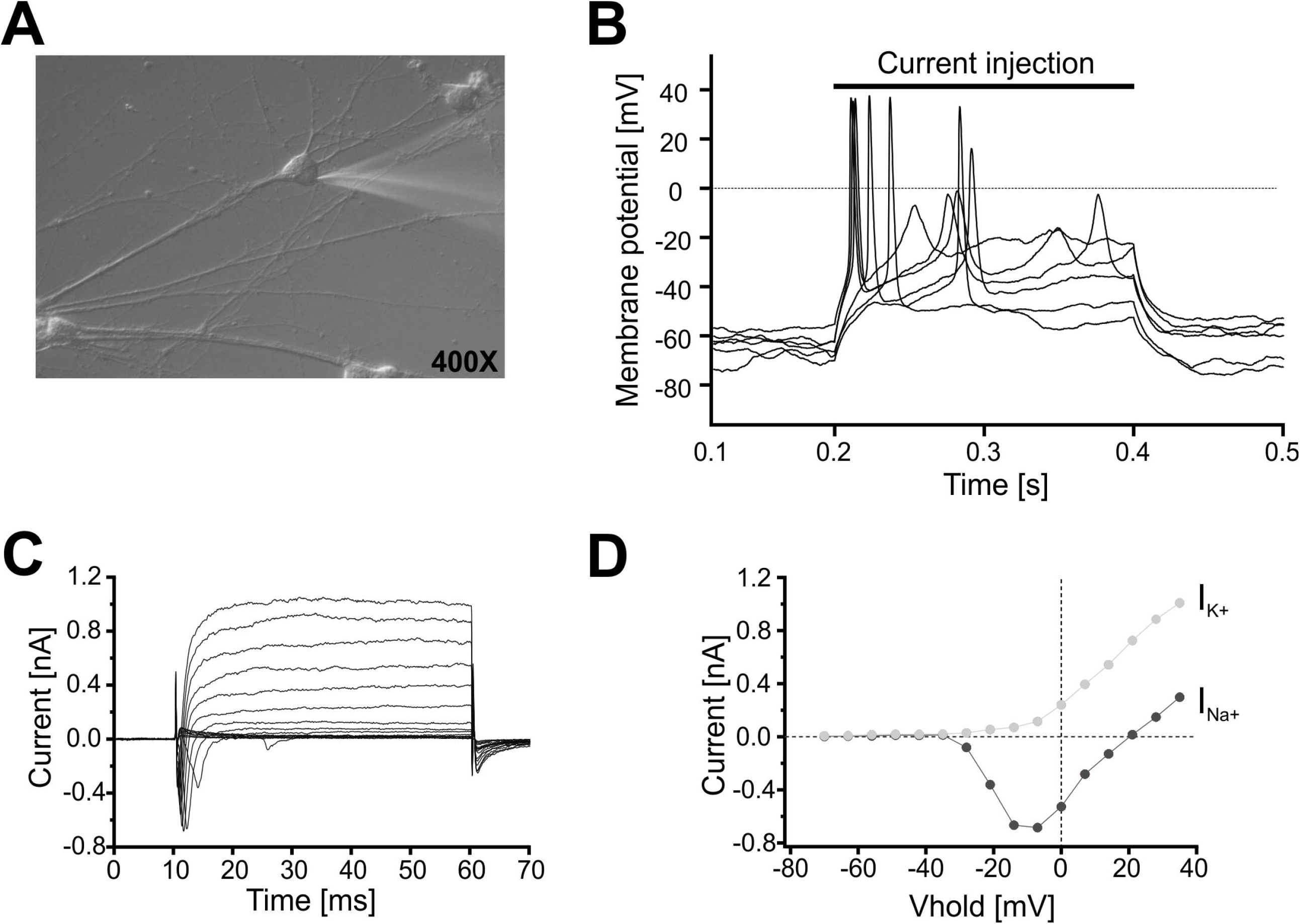
**(A)** Electrophysiological characterization of iPSC-derived neurons. Exemplary differential interference contrast photomicrograph (400x) of a cultured iPSC-derived neuron (DIV 21) from a MDD patient. A patch-pipette is attached to the soma of the cell. **(B)** Whole-cell recording in the current-clamp mode allows recording of the membrane potential. Injection of a supra-threshold depolarizing current induced action potentials. **(C)** Whole-cell recording in the voltage-clamp mode showed voltage-activated transient Na^+^-inward (I_Na+_) and delayed K^+^-outward currents (I_K+_) following a pulse protocol stepping from V_hold_ = -80 mV to more depolarized potentials (−70 to +40 mV). An exemplary current/voltage relationship for voltage-activated Na^+^ (I_Na+_, dark grey filled circles) and K^+^ currents (I_K+_, light grey filled circles) is presented in **(D)**.

**Figure 5.**
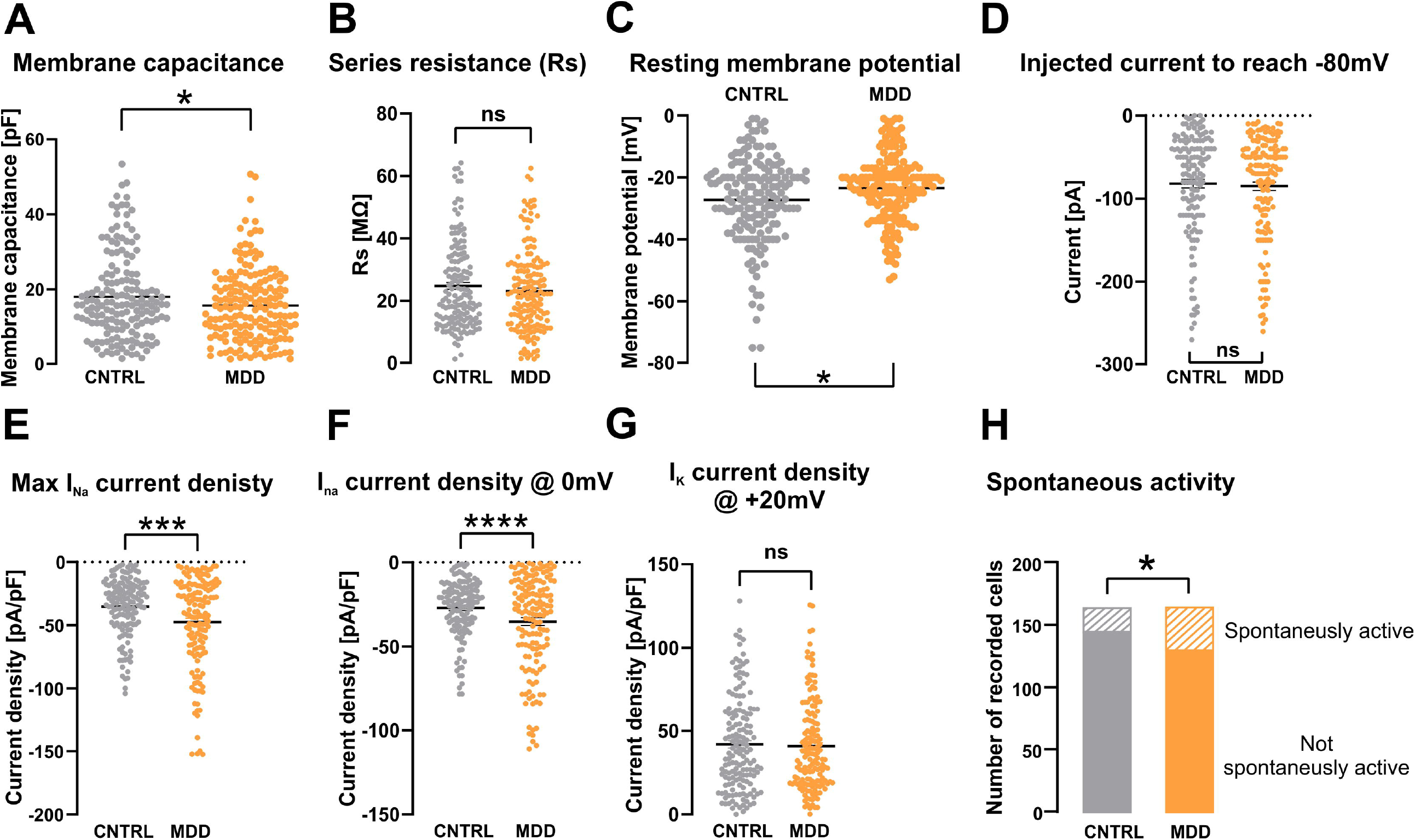
Passive and active electrophysiological properties of iPSC-derived neurons from MDD patients and non-depressed controls. **(A)** Analysis of the membrane capacitance as a measure of the size of the recorded cell revealed a significantly smaller mean value in cells derived from MDD patients. **(B)** Series resistance of iPS-neurons measured in whole-cell voltage-clamp recording. No between-group difference could be detected. **(C)** The resting membrane potential measured in the current-clamp mode was less hyperpolarized in MDD iPS-derived neurons. **(D)** The injected current (in current-clamp experiments) to drive the membrane potential to -80 mV is not different between MDD and CNTRL iPS-neurons. **(E)** Maximum I_Na+_ current density and current density at 0 mV **(F)** is significantly higher in MDD iPS-neurons, whereas I_K+_ current density at +20 mV shows no between-groups difference **(G). (H)** The fraction of spontaneously active iPS-neurons, which fire action potentials at the adjusted membrane potential of -50 mV in a time frame of 30 seconds, was significantly higher in the MDD iPS-neuron group.

## DISCUSSION

In order to identify and characterize molecular pathomechanisms associated with major depression, we established a human cellular model, which is based on dermal fibroblasts derived from skin biopsies of MDD patients and non-depressed controls^6^ and then reprogrammed and differentiated these fibroblasts to iPSCs, NPCs and iPSC-derived neurons.

A reduced value in the pluripotency parameter might be due to contaminating re-differentiated cells (fibroblasts) in the iPSC samples used for the Pluritest. Considering the high proportion of SOX2 and PAX6 positive cells, we decided not to exclude these iPSC lines from our study.

Analysis of the mitochondrial function of reprogrammed human NPCs revealed a reduced function of the OXPHOS in MDD NPCs. These findings indicate a significant mitochondrial alteration, which was already evident in the founder cells (i.e., the fibroblasts) of patients suffering from depression^6^. Other functional parameters of the bioenergetic status of the NPCs, such as MMP or cellular ATP content were not found to be significantly different between the MDD and the control group. A partially altered functional phenotype might in part be explained by an altered metabolic dependence of NPCs. Whereas fibroblasts rely highly on OXPHOS for ATP production, NPCs gain a significant part of their energy via the glycolytic pathway^25, 41^. Thus, potential deficits in their mitochondrial OXPHOS and ETC activity might be compensated by the predominant generation of energy through glycolytic metabolism.

The MDD patients were asked to participate in our study and to donate a skin biopsy at the end of their inpatient stay^6^. At this time, the patients received antidepressant medication and were nearly in remission. Importantly, after several cell divisions of fibroblasts *in vitro*, the confounding variability in these samples on the basis of the subjects’ medication use, should be eliminated^10^. Thus, the alterations in mitochondria function we have found in our study might be associated rather with a trait than with the state of depression. The etiology of depression is associated with genetic (polygenic)^42-44^, as well as with various environmental risk factors such as stressful life events directly influencing physiological processes and mental health of an individual, immediately and in later life. These threats are suggested to affect gene-environment (GxE) interactions and epigenetics by altering chromatin thereby leading to altered molecular patterns observed in depression^45-49^. We observed mitochondrial alterations in fibroblasts of depressed patients^6^ but also in iPSC-derived NPCs reprogrammed from these fibroblasts. So far, our observations can not answer the question, whether this functional phenotype relates to a genetic or epigenetic basis. Although much of the epigenetic memory is erased during the reprogramming process, iPSCs retain a transcriptional memory of the original cells upon reprogramming^50-52^. However, analysis of potential risk variants in the genome of our patient’s cells and the investigation of molecular patterns (e.g., DNA methylation and microRNAs) is an important next step in our search for molecular pathomechanisms of depression.

Today, depression is increasingly viewed as a systemic disease with somatic manifestations outside the brain. Following this hypothesis, we have detected mitochondrial alterations in fibroblasts^6^ and in reprogrammed NPCs of depressed patients. In order to search for neuron-specific disease-related phenotypes and to deepen our understanding of the pathomechanisms involved in depression, we differentiated the NPCs of depressed and non-depressed individuals to iPS-neurons and performed electrophysiological characterization of the biophysical properties by means of whole-cell patch-clamp recording. We recorded from the iPS-neurons after 21 days of differentiation and found that the resting membrane potential was significantly less negative in iPS-neurons derived from MDD patients compared to non-depressed controls. Moreover, consistent with a significantly reduced soma size of NPCs, MDD neurons showed a smaller membrane capacitance, which is indicative for a smaller size or altered geometry of the neuron^53-55^. Intriguingly, iPS-neurons from depressed patients showed a higher Na^+^ current density and an increased fraction of spontaneously active cells *in vitro*. In general, a lower resting membrane potential as well as a smaller cell size could be a result of reduced energy availability in consequence of mitochondrial dysfunction^56^, or due to a delayed neural development, since a hyperpolarized resting membrane potential as well as cell size is associated with neuronal maturation^57, 58^. The increased Na^+^ current density and network activity analyzed as the fraction of spontaneously active neurons when recorded in current-clamp mode is an intriguing observation, which needs deeper investigation. Altered neurite growth and morphology of iPS-neurons derived from depressed patients which were resistant to serotonin reuptake inhibitors have recently been demonstrated and were associated with a reduced expression of the Protocadherin alpha genes in these patients^16^. How this altered activity in iPS-neurons *in vitro* relates to interregional network activity and functional connectivity dynamics *in vivo*^59, 60^, is not known, but an interesting and important objective for future investigations.

## LIMITATIONS

Although we paired MDD patients with non-depressed control subjects of the same sex and a similar age, the study is limited by a high interpersonal variability based on the ages of the subjects within the group, differing lifestyles (e.g. sportive activity) and genetic as well as physiological differences. Moreover, the mental health status of a non-depressed control is based on the self-reported absence of any history of depression in his/her former life. We can not control for a putative load of genetic or environmental risk factors which might have accumulated in the controls. In this line of argumentation, we need to analyse nuclear and mitochondrial DNA for genetic risk factors and further elaborate how much of possible epigenetic patterns could be transmitted from fibroblasts to NPCs and iPS-neurons.

We noticed that the neurons showed quite depolarized RMPs compared to other reports^57^. Since such a low RMP renders voltage-gated sodium channels (VGSCs) mostly steady-state inactivated, we adjusted the membrane potential to about -50 mV by current injection to provide a permissive condition/environment in which AP firing may occur. A depolarized (low) RMP points to a possibly immature developmental state of the induced neurons^57^. In general, neuronal development and electrophysiological properties depend also on the differentiation protocol and substrate (feeder layer, presence of glial cells). It is difficult to decide at which point in time neurons grown *in vitro* can be considered fully mature. Although we found that after 21 days of differentiation, the cells showed the typical almond shaped morphology with an extended meshwork of neurites, generated typical action potentials with overshoots and showed expression of neuronal and synaptic markers, we will use older cultures for future investigations to increase the maturity of our cellular model. However, we observed functional differences between MDD and control neurons after 21 days post differentiation indicating a disease-related phenotype at this rather early state, although depression is not considered a classical neurodevelopmental disorder.

## CONCLUSIONS

We suggest that NPCs and neurons derived from iPSCs of MDD patients can be used to identify and characterize molecular and cellular pathomechanisms of the disease. Our findings indicate that disease-specific alterations observed in fibroblasts are also present in induced-NPCs and might affect the physiology of differentiated iPS-neurons after reprogramming and differentiation. This approach may lead to new insights into the molecular biology and pathophysiology of depression as well as to the discovery of new drug targets and successful therapies.

## ACKNOWLEDGEMENT

The authors would like to thank Richard Warth for providing access to the Seahorse device.

## CONFLICT OF INTEREST

The authors declare no conflict of interest.

## FUNDING

The work has been supported by the Deutsche Forschungsgemeinschaft (DFG, German Research Foundation) project number 422182557 to C.H.W., and GRK2174 to C.H.W., K.K., J.T. and I.C.), the Bavarian State Ministry of Science and the Arts (Bavarian Research Networks ForIPS and ForInter, grants to M.J.R.), and the BMBF (Research Grant No. 01EE1401B to T.C.B. and R.R.).

## Notes

### Competing Interest Statement

The authors have declared no competing interest.

