## Supplement for "Induced neural progenitor cells and iPS-neurons from major depressive disorder patients show altered bioenergetics and electrophysiological properties"

Julian Triebelhorn^1*^, Iseline Cardon^1*^, Kerstin Kuffner^1^, Stefanie Bader^1^, Tatjana Jahner^1^, Katrin Meindl^1^, Tanja Rothhammer-Hampl^2^, Markus J. Riemenschneider^2^, Konstantin Drexler^3^, Mark Berneburg^3^, Caroline Nothdurfter^1^, André Manook^1^, Christoph Brochhausen^4,5^, Thomas C. Baghai^1^, Sven Hilbert^6^, Rainer Rupprecht^1^, Vladimir M. Milenkovic^1^, Christian H. Wetzel^1§^

^1^ Department of Psychiatry and Psychotherapy, University of Regensburg, 93053 Regensburg, Germany

^2^ Department of Neuropathology, Regensburg University Hospital, 93053 Regensburg, Germany

^3^ Department of Dermatology, Regensburg University Hospital, 93053 Regensburg, Germany

^4^ Institute of Pathology, University of Regensburg, 93053 Regensburg, Germany

^5^ Central Biobank of the University of Regensburg and the Regensburg University Hospital, 93053 Regensburg, Germany

^6^ Institute of Educational Research, Faculty of Human Sciences, University of Regensburg, 93053 Regensburg, Germany

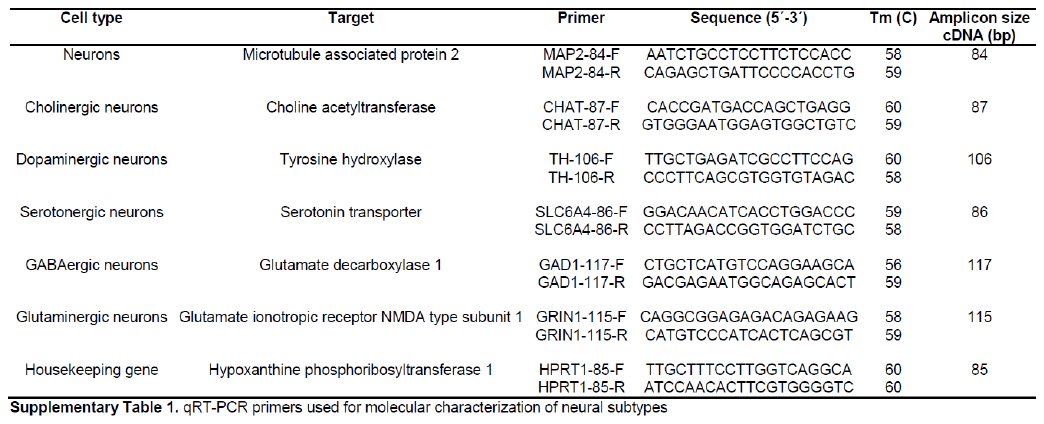

| **ID** | **First diagnose of MDD**  **[year of life]** | **Treatment before hospitalization** | **Treatment during hospitalization** | **Duration of hospitalization [days]** |
| --- | --- | --- | --- | --- |
| MDD1 | 48 | untreated | Mirtazapine | 38 |
| MDD2 | 35 | Citalopram (75 weeks) Opipramol | Venlafaxin | 37 |
| MDD3 | 17 | untreated | Duloxetin, Olanzapine | 107 |
| MDD4 | 35 | Duloxetine, Mirtazapine, Pregabalin, Sertraline, Reboxetine, Olanzapine;  untreated for 2 years before hospitalization | Venlafaxin, Olanzapine | 38 |
| MDD5 | 14 | untreated | Bupropion, Mirtazapine | 62 |
| MDD6 | 21 | untreated | Escitalopram | 43 |
| MDD7 | 23 | untreated | Escitalopram, Mirtazapine | 48 |
| MDD8 | 41 | -2013: Trimipramine, Agomelatine, Mirtazapine, Pregabalin  2013-2015: Amitryptyline, Sertralin  2015-2016: Doxepin | Doxepin | 39 |

**Supplementary Table 2A:** Additional information on medication and hospitalization of MDD patients

| **ID**  **patient** | **Age** | **ID**  **non-depressed CNTRL** | **Age** | **Gender** |
| --- | --- | --- | --- | --- |
| MDD1 | 48 | CON1 | 49 | male |
| MDD2 | 49 | CON2 | 50 | female |
| MDD3 | 19 | CON3 | 25 | male |
| MDD4 | 48 | CON4 | 45 | male |
| MDD5 | 19 | CON5 | 24 | male |
| MDD6 | 24 | CON6 | 25 | female |
| MDD7 | 44 | CON7 | 49 | female |
| MDD8 | 44 | CON8 | 53 | female |

**Supplementary Table 2B:** Information on age and gender

|  | **Hamilton Score** | |
| --- | --- | --- |
| **ID** | **Begin** | **End** |
| MDD1 | 24 | 0 |
| MDD2 | 21 | 8 |
| MDD3 | 24 | 6 |
| MDD4 | 20 | 4 |
| MDD5 | 23 | 12 |
| MDD6 | 23 | 2 |
| MDD7 | 22 | 6 |
| MDD8 | n.a. | n.a. |

**Supplementary Table 2C:** HAM-D ratings at begin and the end of the inpatient stay.

|  | **MAP2 (Ct)** | |
| --- | --- | --- |
| **Group** | **iPSC** | **iNeurons** |
| Con1 | 19.47 | 13.24 |
| Con2 | 18.86 | 13.70 |
| Con3 | 19.02 | 13.78 |
| Con4 | 18.94 | 14.36 |
| Con5 | 20.43 | 14.45 |
| MDD1 | 17.43 | 13.57 |
| MDD2 | 17.68 | 14.27 |
| MDD3 | 18.93 | 13.79 |
| MDD4 | 19.55 | 14.05 |
| MDD5 | 20.60 | 14.72 |
|  | **GAD1 (Ct)** | |
| **Group** | **iPSC** | **iNeurons** |
| Con1 | 24.71 | 16.86 |
| Con2 | 24.96 | 17.70 |
| Con3 | 25.02 | 16.52 |
| Con4 | 25.03 | 15.44 |
| Con5 | 25.18 | 17.30 |
| MDD1 | 24.11 | 18.81 |
| MDD2 | 22.89 | 18.15 |
| MDD3 | 24.72 | 17.58 |
| MDD4 | 24.12 | 15.13 |
| MDD5 | 25.37 | 18.27 |
|  | **GRIN1 (Ct)** | |
| **Group** | **iPSC** | **iNeurons** |
| Con1 | 26.49 | 18.79 |
| Con2 | 26.97 | 17.08 |
| Con3 | 28.74 | 19.28 |
| Con4 | 27.23 | 16.35 |
| Con5 | 28.29 | 19.26 |
| MDD1 | 26.50 | 19.31 |
| MDD2 | 26.43 | 19.08 |
| MDD3 | 28.62 | 18.93 |
| MDD4 | 29.06 | 17.66 |
| MDD5 | 28.42 | 18.30 |
|  | **SLC6A4 (Ct)** | |
| **Group** | **iPSC** | **iNeurons** |
| Con1 | 23.93 | 20.55 |
| Con2 | 24.35 | 20.98 |
| Con3 | 25.17 | 22.02 |
| Con4 | 24.86 | 23.13 |
| Con5 | 24.32 | 20.63 |
| MDD1 | 24.41 | 21.7 |
| MDD2 | 25.11 | 22.89 |
| MDD3 | 24.45 | 21.01 |
| MDD4 | 25.4 | 21.72 |
| MDD5 | 24.1 | 22.99 |
|  | **TH (Ct)** | |
| **Group** | **iPSC** | **iNeurons** |
| Con1 | 25.89 | 24.02 |
| Con2 | 28.7 | 22.72 |
| Con3 | 27.72 | 18.34 |
| Con4 | 27.11 | 21.57 |
| Con5 | 27.82 | 20.18 |
| MDD1 | 28.34 | 24.95 |
| MDD2 | 28.11 | 22.92 |
| MDD3 | 27.84 | 20.7 |
| MDD4 | 26.63 | 22.43 |
| MDD5 | 27.21 | 18.65 |
|  | **CHAT (Ct)** | |
| **Group** | **iPSC** | **iNeurons** |
| Con1 | 25.37 | 20.8 |
| Con2 | 25.64 | 21.83 |
| Con3 | 26.4 | 20.86 |
| Con4 | 25.96 | 22.07 |
| Con5 | 26.73 | 20.73 |
| MDD1 | 28.68 | 21.98 |
| MDD2 | 26.83 | 23.17 |
| MDD3 | 28.62 | 20.02 |
| MDD4 | 27.19 | 20.68 |
| MDD5 | 25.28 | 21.82 |

**Supplementary Table 3.** List of Ct values of RT-PCR reactions used for generation of gene expression heat maps.

| **Code** | **ID** | **Neurons recorded**  **(total)** | **Used in statistical analysis** |
| --- | --- | --- | --- |
| K | MDD3 | 26 | 24 |
| L | Con3 | 19 | 19 |
| Q | Con7 | 35 | 30 |
| M | Con5 | 29 | 23 |
| N | MDD2 | 30 | 21 |
| O | Con2 | 23 | 17 |
| P | MDD5 | 23 | 12 |
| V | Con4 | 21 | 11 |
| U | MDD4 | 29 | 20 |
| F | MDD1 | 32 | 18 |
| E | Con1 | 29 | 12 |
| Y | Con8 | 28 | 21 |
| Z | MDD8 | 30 | 21 |
| R | MDD7 | 30 | 26 |
| A | Con6 | 31 | 29 |
| B | MDD6 | 31 | 29 |
| Total |  | 446 | 333 |

**Supplementary Table:** Additional information for patch clamp experiments

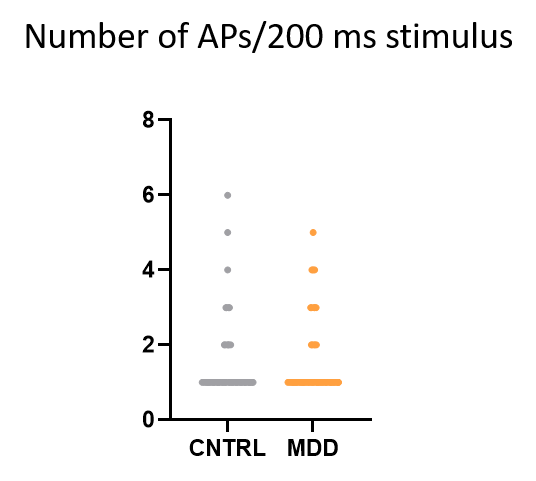

**Supplementary Figure**: Number of APs during a depolarizing stimulus (200 ms). No between-group difference in AP firing activity could be detected.
